# Semisynthesis reveals Apoptin as a tumour-selective protein prodrug that causes cytoskeletal collapse

**DOI:** 10.1101/2022.11.23.517692

**Authors:** Jasmine Wyatt, Mahvash Tavassoli, Manuel M. Müller

**Affiliations:** Department of Molecular Oncology, King’s College London, Guy’s Hospital Campus, Hodgkin Building, London SE1 1UL, UK; Department of Chemistry, King’s College London, Britannia House, 7 Trinity Street, London, SE1 1DB, UK

## Abstract

Apoptin is a small viral protein capable of inducing cell death selectively in cancer cells. Despite its potential as an anticancer agent, relatively little is known about its mechanism of toxicity and cancer-selectivity. Previous experiments suggest that cancer-selective phosphorylation modulates Apoptin toxicity, although a lack of chemical tools has hampered the dissection of underlying mechanisms. Here, we describe structure-function studies with site-specifically phosphorylated Apoptin (Apoptin-T108ph) in living cells which revealed that Thr108 phosphorylation is the selectivity switch for Apoptin toxicity. Mechanistic investigations link T108ph to actin binding, cytoskeletal disruption and downstream inhibition of anoikis-resistance as well as cancer cell invasion. These results establish Apoptin as a protein pro-drug, selectively activated in cancer cells by phosphorylation, which disrupts the cytoskeleton and promotes cell death. We anticipate that this mechanism provides a framework for the design of next generation anticancer proteins with enhanced selectivity and potency.

## INTRODUCTION

One of the major challenges of current cancer therapeutics is to improve their cancer-selectivity. Most existing chemotherapies target functions common to both tumour and healthy cells, limiting their therapeutic efficacy and resulting in adverse side effects. Apoptin was originally identified as the apoptosis-inducing VP3 protein from the chicken anaemia virus (CAV), the first member of the Gyrovirus genus ^1^. This fascinating protein has an intrinsic ability to selectively kill a wide variety of tumour cells with little cytotoxicity in normal cells ^2,3^. Studying its mechanism of action will inform on strategies for achieving cancer-selective toxicity.

Apoptin is a small (14 kDa), intrinsically disordered protein rich in proline, serine, threonine and basic amino acids. It contains a bipartite nuclear localisation signal (NLS) and a nuclear export sequence (NES), key in facilitating the transport of the protein between the nucleus and cytoplasm (Fig. 1a) ^4^. The mechanisms of activation of Apoptin and consequent cell death-inducing pathways in cancer cells are largely unknown ^5^. Post-translational modifications (PTMs) are important molecular switches for regulating protein function, subcellular localisation and the protein’s interactome. Apoptin is decorated with several PTMs, particularly in the form of phosphorylation. Threonine-108 phosphorylation (T108ph) is known to be tumour-specific and associated with the accumulation of the protein in the nucleus of cancer cells via the inactivation of the adjacent NES ^6,7^. This PTM occurs by tumour-specific kinases such as an isozyme of protein kinase C (PKCβ), earlier discovered to cooperate with Apoptin in colorectal cancer cells ^7^. Prior mutagenesis studies have yielded some valuable insight into the importance of the phospho modification within Apoptin’s C-terminus, although with inconsistent results. Mutation of T108 in several studies only partially affected the toxic activity in tumour cells ^8,9^. However, it has been noted that a T108A mutation causes T106 and T107 to become opportunistically phosphorylated as an alternative, allowing this mutant to maintain tumour-selective death ^10^. Additionally, the introduction of a negative charge to mimic phosphorylation via site-directed mutagenesis (T108E) enabled Apoptin to accumulate in the nucleus and force the sensitisation of healthy cells ^6,11^. Nonetheless, phospho-mimics have been shown to often fail to capture the complete function of Thr phosphorylation due to discrepancies in overall charge, size and geometry of the residue ^12^. Moreover, the mechanism by which apoptin phosphorylation leads to cell death is unknown.

**Figure 1:**
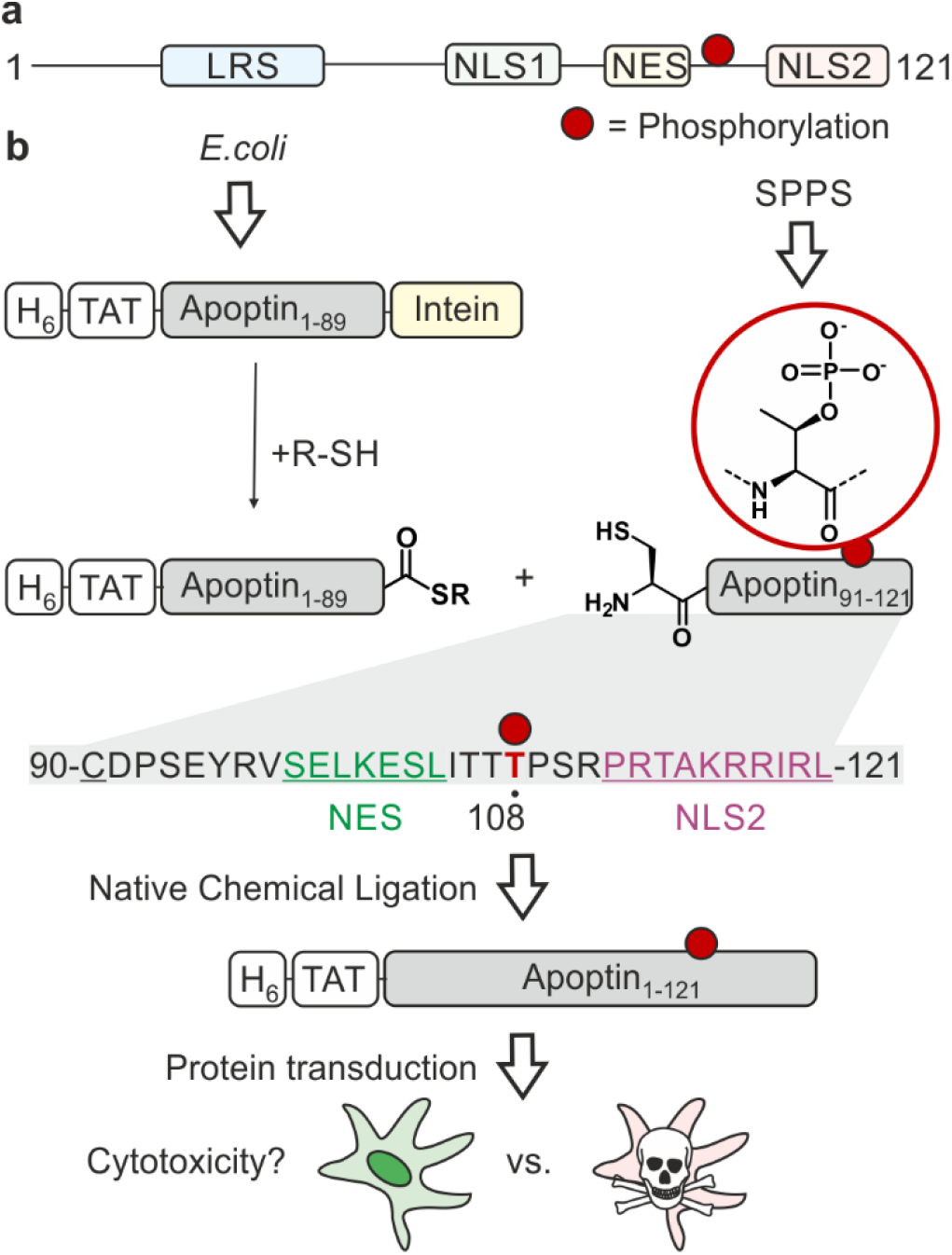
Strategy for the semisynthesis and application of Apoptin. a) Domain architecture of Apoptin. LRS = Leucine rich sequence; NLS = Nuclear localisation sequence; NES = Nuclear export sequence. Phosphorylation at threonine 108 (T108ph) is indicated with solid red circle. b) Semisynthesis strategy to access site-specifically phosphorylated Apoptin. The N-terminus of Apoptin is made recombinantly with a hexahistidine tag (H_6_) fused to the TAT cell-penetrating peptide. The desired C-terminal α-thioester is obtained via Intein technology. The N-terminal peptide is synthesised chemically with an N-terminal cysteine (Cys90, underlined) to enable native chemical ligation (sequence shown below with NES and NLS2 in green and purple, respectively. Site of modification shown in red). The Apoptin building blocks can then be joined together using native chemical ligation yielding native Apoptin which will be deployed to probe the effect of T108ph in living cells upon protein transduction via the TAT peptide.

Therefore, we sought new chemistry-driven approaches to interrogate the role of site-specific phosphorylation of Apoptin in toxicity and tumour selectivity. We developed a protein semisynthesis strategy to generate site-specifically phosphorylated Apoptin (Apoptin-T108ph) furnished with a cell-penetrating peptide, enabling us to directly measure the functional consequences of this modification in living cells. Toxicity studies with semisynthetic apoptin in cancer-derived and healthy cell lines demonstrate that Apoptin acts as a protein pro-drug and is activated by C-terminal phosphorylation into a non-specific protein toxin. Pulldowns with Apoptin-derived (phospho-)peptides revealed that phosphorylation enhances Actin binding, suggesting a new mechanism of action. Follow-on microscopy and functional studies confirm interactions with cytoskeletal components and provide evidence that Apoptin perturbs cytoskeleton structure and function. This study thus provides a mechanistic framework for how Apoptin acts as an anti-tumour agent and suggests a novel strategy for the design of cancer-specific protein prodrugs that target the cytoskeleton in a phospho-dependent manner.

## RESULTS & DISCUSSION

### Semisynthetis of (phospho-)Apoptin

Synthetic phospho-proteins are powerful reagents to explore the role of phosphorylation in modulating protein structure and function ^12^. In conjunction with protein transduction methods ^13,14^, it is possible to deploy chemically tailored proteins for functional studies in living cells ^15–19^. We therefore developed a protein semisynthesis strategy ^20^ to generate site-specifically modified Apoptin. This modular approach harnesses the combination of synthetic peptides carrying the defined PTM and recombinant protein fragments making up the bulk of the unmodified protein. These building blocks are then joined together by native chemical ligation (NCL) yielding a native peptide bond. NCL requires chemically compatible reaction handles *i*.*e*. a *C*-terminal α-thioester and an *N*-terminal cysteine residue ^21^. The *C*-terminus of Apoptin contains the PTM sites of interest, and features a native cysteine residue at position 90. Hence, we decided to chemically synthesise the region 90-121 preserving the natural amino acid sequence of Apoptin and enabling the introduction of phospho-amino acids. The remainder of the protein could then be expressed in *E*.*coli* as a fusion with the MxeGyrA intein to access a *C*-terminal α-thioester (Fig. 1b).

We began by synthesizing an unmodified Apoptin peptide encompassing residues 90-121 (Apoptin_90-121_) on solid phase. Semi-automated solid-phase peptide synthesis (SPPS) was exploited using carbodiimide/Oxyma couplings and *N*-α-Fmoc-protected amino acids. Following cleavage and purification by reverse phase (RP)-HPLC, 4 mg of the peptide (isolated yield ≈5%) with an *N*-terminal Cys90 was obtained (Fig. S1a).

To access the N-terminal fragment of Apoptin with a *C*-terminal α-thioester, we opted for an Intein approach. This method utilizes the mechanism of Intein self-splicing in order to generate recombinant proteins with a C-terminal α-thioester ^22^ (Fig. S2). Briefly, truncated Apoptin (residues 1−89, Apoptin_1-89_) was expressed in fusion with the *Mxe* GyrA mini-intein (Fig. S2e). A hexahistidine tag was retained at the N-terminus to facilitate comparison with previous variants produced by recombinant means and an HA-tag and TAT sequence were included to facilitate detection and for cell-penetration ^23–25^, respectively. The construct was produced in *E. coli* via auto-induction and purified from the resulting inclusion bodies. Intein thiolysis was achieved with the addition of 0.1 M MESNa to obtain the reactive thioester required for ligation (Apoptin_1-89_-SR) which was subsequently isolated with a final yield of ≈25% by RP-HPLC (Fig. S2b,c,d,f).

Chemoselective ligations of Apoptin_1-89_-SR and Apoptin_90-121_ in GdmCl in the presence of 50 mM mercaptophenylacetic acid (MPAA) proceeded smoothly over the course of 2 hours, as judged by RP-HPLC and SDS-PAGE (Fig. 2a-c). Within 2 h, the peak corresponding to the Apoptin-thioester had decreased, with simultaneous emergence of a new peak at 17.6 min retention time (Fig. 2b). Likewise, a gel shift from ∼15 kDa to ∼20 kDa is observed, consistent with ligation to full-length Apoptin (Fig. 2c). After 2 h, considerable hydrolysis of the protein thioester occurred and no major increase in product was observed (Fig. S3). Upon RP-HPLC purification, we obtained 1 mg of full-length Apoptin (Apoptin-WT*, Fig. 2d) which was confirmed with high-resolution mass spectrometry (HRMS, expected mass 18562.9 Da, experimental mass 18561.6 Da) (Fig. 2e). The isolated yield (≈ 7.5%) was lower than expected due to the difficulty in separating the full-length protein from Apoptin_1-89_-SR.

**Figure 2:**
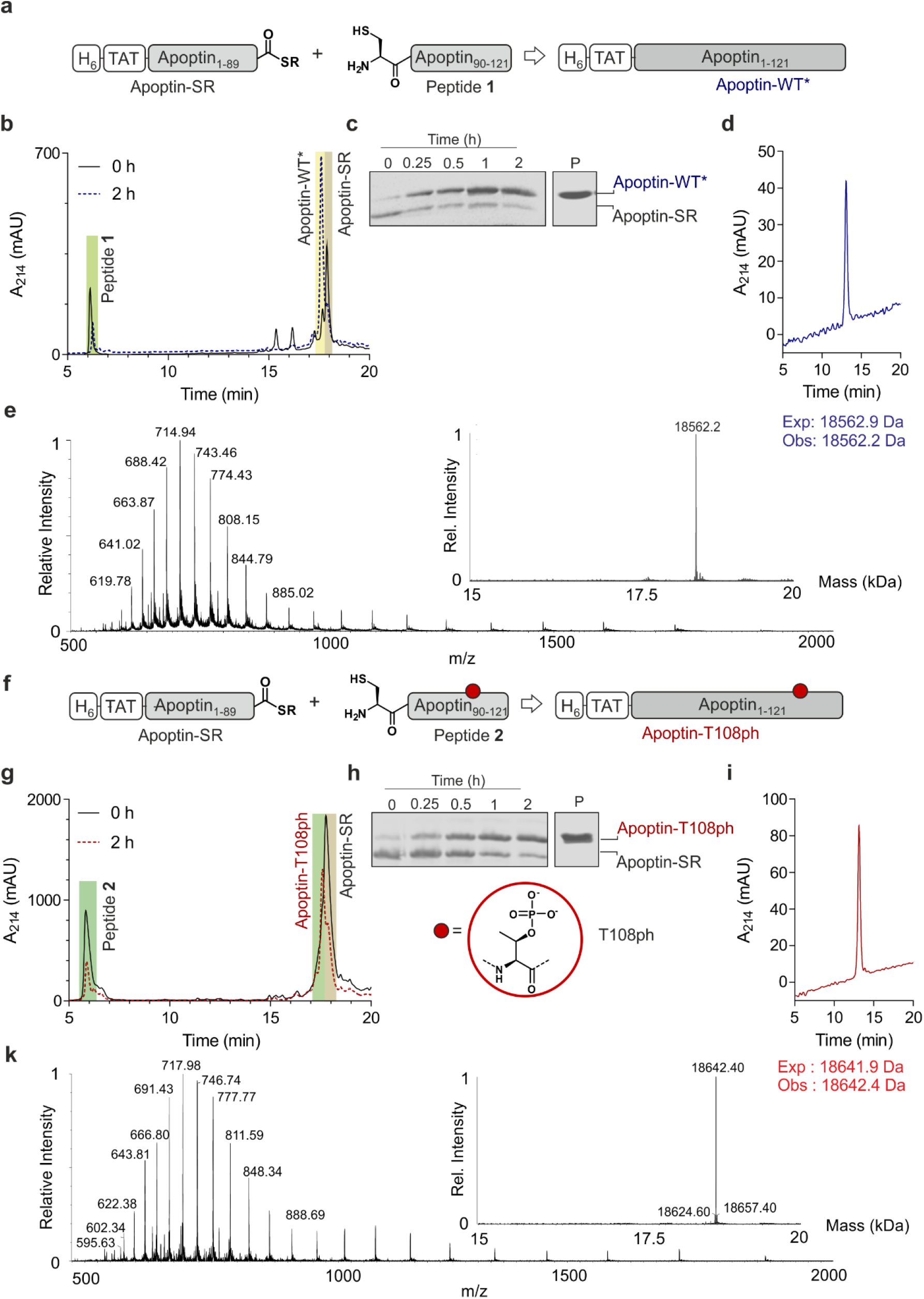
Native chemical ligation of Apoptin. a) Schematic representation of the ligation reaction to synthesise Apoptin-WT*. b) Time course of the Apoptin-WT* ligation reaction monitored by RP-HPLC. c) Time course of the unmodified Apoptin ligation reaction monitored by SDS-PAGE, P: isolated and renatured Apoptin-WT*_ified_. d) RP-HPLC of isolated Apoptin-WT*. e) Mass spectrometry analysis of final isolated Apoptin-WT* protein (expected mass 18562.9 Da, observed mass 18561.6 Da). f) Schematic representation of the ligation reaction to synthesize modified Apoptin-T108ph. g) Time course of the modified Apoptin-T108ph ligation reaction monitored by RP-HPLC. h) Time course of the modified Apoptin-T108ph ligation reaction monitored by SDS-PAGE, P: isolated and renatured modified Apoptin-T108ph. i) RP-HPLC of isolated modified Apoptin-T108ph. k) Mass spectrometry of final isolated Apoptin-T108ph (expected mass 18641.9 Da, observed mass 18642.4 Da.

Encouraged by the successful synthesis of full-length Apoptin, we next targeted the synthesis of the phosphorylated derivative Apoptin-T108ph. As before, the modified T108ph peptide 2 was synthesised using SPPS with incorporation of Fmoc-Thr(PO(OBzl)OH)-OH using HATU couplings. The phospho-peptide was isolated after two rounds of purification in sufficient amounts (4mg, ≈ 0.5% isolated yield) and purity to proceed with ligation to Apoptin_1-89_-SR (Fig. S1c,d). The ligation proceeded rapidly (Fig. 2f,g,h); after 2h, the reaction was quenched and full-length Apoptin-T108ph was isolated (Fig. 2i). HRMS analysis (Expected mass 18641.9 Da, experimental mass 18642.4 Da) confirmed that this species contains a single phosphoryl group (Fig. 2k).

Finally, we set out to renature our semisynthetic derivatives into biologically active Apoptin, which is thought to be intrinsically disordered ^26^. To optimize the renaturation procedures, we produced full-length recombinant Apoptin bearing an N-terminal hexahistidine tag and TAT cell-penetrating peptide in *E. coli* and purified the protein directly from inclusion bodies (Apoptin-WT). We then renatured Apoptin-WT from 6 M GdmCl into 25 mM MES, 100 mM NaCl, 2 mM MgCl_2_ via a one-step dialysis procedure. RP-HPLC, MS and SDS-PAGE confirmed the identity and purity of the final Apoptin-WT preparations (Fig. S4) Once this procedure was optimised, we repeated this for the unmodified and T108ph semisynthetic Apoptin products (Fig. 2d,i shows isolated and renatured semisynthetic material).

### Apoptin is a pro-drug activated by phosphorylation

To ensure that our unmodified proteins retain cancer-specific toxicity as reported in the literature ^3,27^, we first tested the Apoptin-WT in a panel of cancerous and healthy cell-lines. Using variable protein doses, we measured cell death after 24 hours using an MTT assay. The results showed that Apoptin-WT is toxic in two different tumour cell-lines (SAOS-2 and HSC-3) in a dose-dependent manner. Importantly and in agreement with prior work, healthy cell-lines (HGF and NHDF) were resistant to Apoptin treatment (Fig. 3a). Semisynthetic Apoptin-WT* showed similar toxicity when compared to the Apoptin-WT (Fig. 3b), highlighting the overall feasibility of our combined semisynthesis/protein transduction approach.

**Figure 3:**
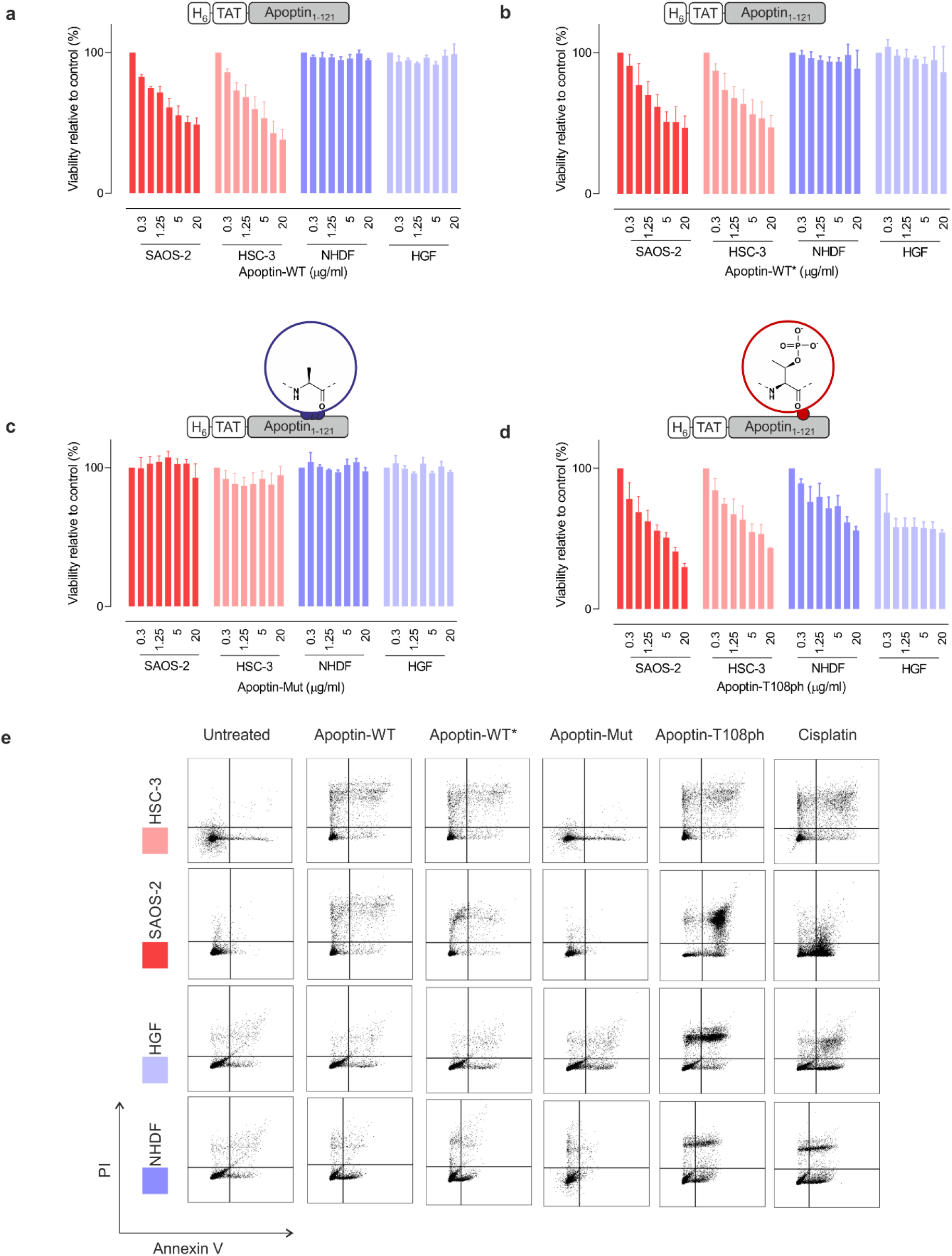
Apoptin toxicity is activated by phosphorylation at Thr108. MTT assay in cancer cells (red) and healthy cells (blue) when treated for 24 h with varying doses of (a) Apoptin-WT; (b) Apoptin-WT*; (c) Apoptin-Mutant; and (d) Apoptin-T108ph (n=3 ± SEM). (e) FACS with Annexin/PI staining for cancer cells (red) and healthy cells (blue) treated with different Apoptin derivatives (5 μg/mL) and Cisplatin (positive control) for 24 h. Full analysis shown in supplementary Figure S6.

Previous research has shown that mutation of T108 to alanine abolishes apoptotic behaviour but other research postulated that mutation of this residue only promotes phosphorylation of adjacent threonines (106-107) instead, allowing Apoptin induced cell death to continue to occur ^6,10^. To preclude phosphorylation in this region, we mutated all three threonine residues to alanine (T106-108>A) and produced the resulting variant (Apoptin-Mut, Fig. S5). We demonstrated that this derivative has very limited toxicity in all cell-lines tested, indicating that phosphorylation of T108 is required for Apoptin-induced toxicity (Fig. 3b). Next, we measured the cytotoxicity of C-terminally phosphorylated Apoptin-T108ph. Remarkably, healthy cell-lines – which were resistant to unmodified Apoptin – are sensitive to treatment with Apoptin-T108ph (Fig. 3d). FACS analysis with AnnexinV (FITC)-PI double staining qualitatively confirmed these results (Fig. 3e and S6). This result directly demonstrates that phosphorylation at T108 is a key driver for Apoptin toxicity and thus represents the cancer-specific kill-switch of this protein.

### Phosphorylation of Apoptin promotes an interaction with the Actin cytoskeleton

To gain insight into the mechanism of action of activated Apoptin-T108ph, we surveyed binding to a range of interaction partners previously described in the literature. For this purpose, we synthesised and purified biotinylated C-terminal Apoptin peptides with and without the T108ph modification (Fig. S7). To examine the effect of phosphorylation on the Apoptin-associated cellular proteins, we immobilised the newly synthesised peptides on streptavidin beads, incubated them with cell extracts from HSC-3 and analysed bound proteins by western blotting.

We confirmed the previously reported interaction between Apoptin and the Anaphase-promoting complex/cyclosome (APC/C) subunit APC1 ^28^. However, we noticed that this association was in fact hindered when Apoptin is phosphorylated at position T108 (Fig. S8a,b). Moreover, we show for the first time that Apoptin can physically interact with the nuclear export protein CRM-1, likely via the postulated NES ^29^ between position 97-105 (Fig. S8a,b). Contrary to previous hypotheses ^29^, the NES is not completely abolished by phosphorylation, at least under the conditions of our pull-down assay. No binding to nuclear beta-importin was observed (Fig. S8c), presumably because our biotinylated peptides only encompass the second fragment of the bipartite NLS.

Previous work showed that flag-tagged Apoptin co-immunoprecipitated with alpha-tubulin, beta-tubulin and beta-Actin ^28^. We therefore tested the impact of phosphorylation on these interactions. Interestingly, T108ph-containing peptides bound more readily to Actin than unmodified counterparts, suggesting Actin as a potential target for phospho-Apoptin toxicity (Fig. 4a,b). No interaction with alpha-tubulin was detected, suggesting that full-length Apoptin may be required for this interaction (Fig. 4a,b). To further validate this finding, cancer cells were treated with full-length recombinant Apoptin and analysed for distribution by immunofluorescence. The cellular distribution of transduced Apoptin-WT resembled Actin stress fibres and showed a high degree of co-localisation with F-Actin, further reinforcing a relationship between Apoptin and the cytoskeleton (Fig. 4c,d).

**Figure 4:**
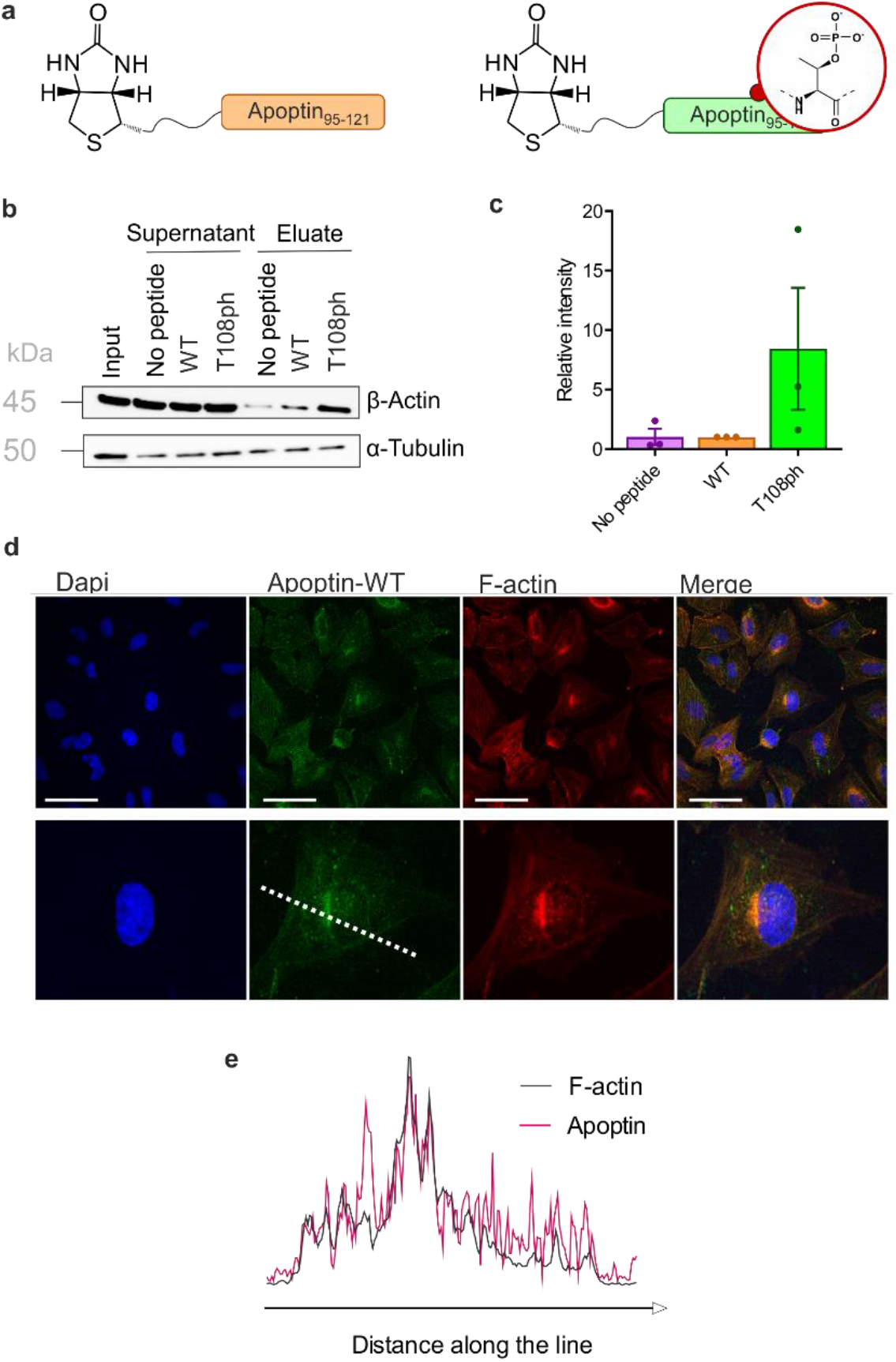
T108 phosphorylation promotes interaction with the Actin cytoskeleton. a) Schematic of biotinylated C-terminal Apoptin peptides used in this study. B) Western blot analysis of pull-downs with wild-type and T108ph Apoptin peptides. b) Quantification of β-Actin pull-downs showing that β-Actin preferentially bound to the T108ph-modified Apoptin peptide (n=3 ± SEM). Full replicates are shown in Supplementary Fig. 7c) Immuno-fluorescence of SAOS-2 cells treated with Apoptin and fixed and stained with anti-HA or TRITC-phalloidin. Scale bars 100 μm. The image below shows a single cell zoomed in to illustrate a build-up of Apoptin in peri-nuclear regions and a colocalisation of Apoptin and F-Actin. d) Line scan of this cell was performed by drawing a single line through the middle of the cell (dotted white line in c).

### Apoptin causes cytoskeletal collapse in tumour cells

This novel finding showing that Apoptin phosphorylation results in increased interactions with the cytoskeleton led us to investigate the effect of Apoptin treatment on cancer cell architecture. HSC-3 cells were treated with sublethal doses (0.5 μg/mL) of Apoptin to ensure limited effects on cellular viability and proliferation before being fixed and stained for E-cadherin and F-Actin (Fig. 5a). Densely cultured HSC-3 cells have a marginal Actin bundle with established adhesion junctions aligned as a continuous belt along the cell-cell boundaries. We found that treatment with Apoptin-WT resulted in clear alterations in the Actin cytoskeleton and the appearance of fewer cell-cell contacts. Importantly, this observation was not seen in the samples treated with Apoptin-Mut which cannot be phosphorylated at position 108. This pattern is tumour specific as HGF cells treated with recombinant or mutant Apoptin do not display any obvious changes in the Actin cytoskeleton (Fig. S9). Moreover, we ruled out that this effect was due to a direct interaction between Apoptin and E-cadherin because C-terminal Apoptin peptides were unable to bind to E-cadherin (Fig. S8c). To accurately quantify observed changes in F-Actin, cells were permeabilised and stained with TRITC-Phalloidin for analysis by flow cytometry. HSC-3 cells demonstrated a loss of filamentous Actin when treated with sublethal doses of recombinant Apoptin, strengthening the immunofluorescence results (Fig. 5b).

**Figure 5:**
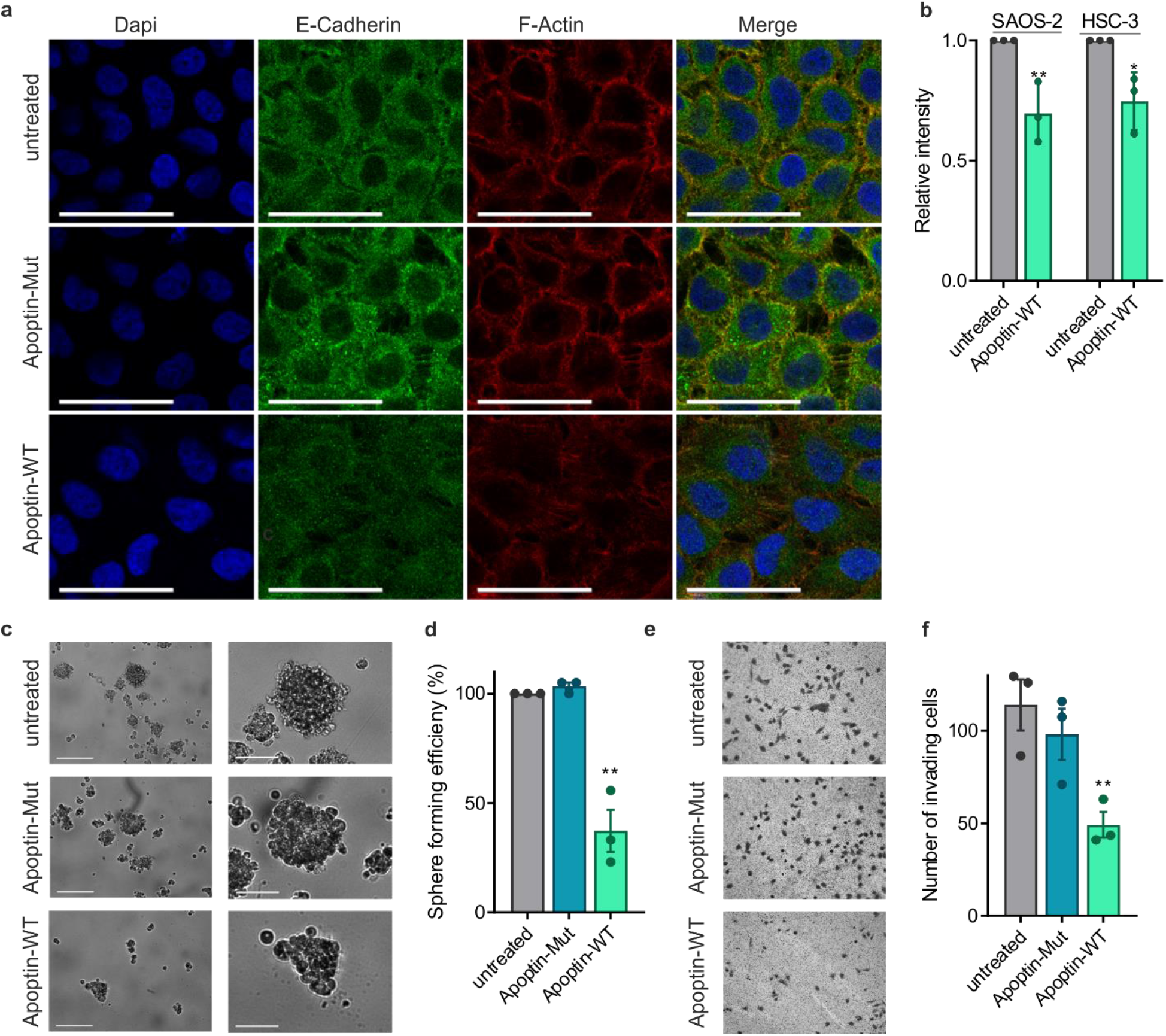
Apoptin can inhibit cancer cell metastasis. a) Immuno-fluorescence images of HSC-3 cells treated with sublethal doses (0.5 μg/mL) of Apoptin-WT or Apoptin-Mut and stained for E-Cadherin and F-Actin. Cells were imaged at 60x magnification, scale bar=40 μm. b)FACS analysis of total F-Actin in HSC-3 cells treated with sublethal doses (0.5 μg/mL) Apoptin-WT. c) Representative image of cells grown in low-adherent conditions for 7 days after treatment with the relevant protein. Images below show the biggest spheroid formed on each plate. Scale bars 200 μm (top) and 80 μm (bottom). d) Quantification of spheroid efficiency. e) Representative images of invading cells in each condition. (f) Quantification of number of invading cells (n=3, error bars ± SEM *p<0.05 **p<0.01.).

Cancer metastasis is a multi-step process that is ultimately driven by localised polymerisation of Actin filaments. Since Apoptin has a natural ability to disrupt F-Actin and cancer cell architecture, we next set out to assess anoikis resistance and invasion in cancer cells after Apoptin treatment. HSC-3 cells are intrinsically anoikis resistant and are able to form several multicellular spheroids when cultured in low-adherent conditions (Fig. 5c,d). Treatment with recombinant Apoptin significantly reduced number of spheroids (p<0.01), while mutant Apoptin had little effect on spheroid formation. In addition, Matrigel assays were used as a measure of cell invasion. HSC-3 cells were found to be highly invasive, but the number of invading cells was significantly decreased upon treatment with recombinant wild-type Apoptin treatment (p<0.01), but not with mutant Apoptin. Collectively, these results suggest that, upon activation in cancer cells by phosphorylation, Apoptin binds to and interferes with cytoskeletal components, contributing to cytoskeletal collapse.

## CONCLUSION

Apoptin has gained some attention as a promising cancer-specific drug, however its therapeutic potential has not progressed due to the lack of understanding of its mechanism, precluding the engineering of next-generation variants. The lack of insight into Apoptin is due to its complex mechanism of action involving post-translational modifications, and, accordingly, difficulties in obtaining defined forms of the protein. To bypass these issues, we developed a protein semisynthesis strategy, allowing us to establish the role of Apoptin phosphorylation in its toxicity and cancer selectivity. Although the high number of consecutive β-branched residues present in Apoptin’s C-terminus negatively affected the yield of synthetic (phospho-)peptides, sufficient quantities of peptides were obtained to complete the semi-synthesis and perform toxicity studies in live cells. Our approach will furthermore enable the introduction of spectroscopic probes and crosslinkers into the C-terminal region of Apoptin, further facilitating mechanistic studies of its activity and selectivity.

Previous research indicated an important role for Thr108 phosphorylation in apoptin function^6,7^, but how this modification contributes to toxicity has remained unknown. By deploying semisynthetic full-length Apoptin-T108ph we proved that phosphorylation acts as the switch for toxicity and provided new evidence for its mechanism of action. The observation that healthy cells, previously resistant to Apoptin, are susceptible to the semisynthetic protein with the addition of T108ph demonstrates that Apoptin itself is not toxic. Instead, phosphorylation – installed by kinases hyperactive in cancer – activates its toxicity, establishing Apoptin as a cancer-selective prodrug. This mechanism of activation is distinct from traditional prodrugs which typically consist of an anticancer drug fused to a cleavable linker, activated only once the linker is split by constituents in the tumour environment. Linker cleavage can be promoted by physiochemical features such as hypoxia ^30^, low pH ^31^ reactive oxygen species (ROS) ^32^ or physical triggers including photoactivation ^33^. Prodrug activation may exploit secreted overexpressed tumour proteins such as extracellular matrix metalloproteinases ^34^ and aminopeptidases ^35^ or intracellular enzymes such as DT-diaphorase ^36^, carboxylesterases ^37^ and β-galactosidase ^38^. The mode of Apoptin prodrug activation also complements previously explored protein prodrug design strategies, operating via selective release of toxic proteins upon exposure to disease-state specific enzymatic activities ^39^. Our work thus provides a conceptual basis for the design of exogenously added protein prodrugs that can be directly and selectively activated (in this case by phosphorylation) by endogenous enzymes. Given that this mechanism is compatible with other means of achieving tumour selectivity, Apoptin could be used as a starting point to construct ultra-selective anticancer proteins.

In addition, our strategy identifies a possible mechanism of action of Apoptin through phospho-dependent protein-protein interactions. We showed that Apoptin phosphopeptides interact with beta-Actin providing a direct link between apoptin activation, the cytoskeleton and cell death. Previous research has noted that Apoptin, when over-expressed in cancer cell lines, localised to the cytoplasm initially where immunofluorescence revealed a filamentous staining resembling Actin stress fibres ^23^. Interestingly, other viral proteins with similar characteristics to Apoptin have also been suggested to act via cytoskeletal remodelling (reviewed in ^5^). Our data indicates that the interaction between Apoptin and the Actin cytoskeleton is phosphorylation-dependent and leads to a collapse of filamentous architecture. Consequently, we showed that in tumour cells, Apoptin treatment limits invasion and anchorage abilities, characteristics that are key for cancer progression. With invasion and metastasis accounting for more than 90% of mortality in solid cancers, there is a need for the development of anti-invasion/anti-metastasis drugs that could be used in combination with antiproliferative therapy ^40,41^. Based on our results, Apoptin is a promising candidate for such a strategy and could potentially function as a migrastatic.

Collectively, our results demonstrate the origin of cancer selectivity and a potential mechanism of action of the viral anticancer protein Apoptin. Our work therefore provides an avenue for designing new cancer-selective protein prodrugs which are also capable of inhibiting invasion and metastasis.

## METHODS

### Materials

Saos-2 (human osteosarcoma cell-line) was a gift from Gerry Melino, University of Leicester. HSC-3 (human head and neck cancer cell line) was a gift from Kazuya Tominaga, Department of Oral Pathology, Osaka Dental University. HGF (early passage secondary culture of normal human gingival fibroblasts) was a gift from Francis Hughes, Dental Institute, Kings College London and NHDF (early passage secondary culture of normal human dermal fibroblasts) was a gift from Tanya Shaw, Division of Immunology, Infection & Inflammatory Disease, King’s College London. All cell lines were maintained in DMEM, supplemented with 10% foetal bovine serum (FBS) and 20μg/ml penicillin, 10μg/ml streptomycin and 1mM sodium pyruvate (All Sigma).

Peptide resins, Fmoc-L-amino acids, the monoprotected phosphothreonine building block (Fmoc-Thr(PO(OBzl)OH)-OH), Hexafluorophosphate Azabenzotriazole Tetramethyl Uronium (HATU), Oxyma and K-Oxyma were purchased from Novabiochem. Diisopropylethylamine (DIEA), piperidine, phenol, Tris(2-carboxyethyl)phosphine hydrochloride (TCEP) were purchased by Merck Sigma-Aldrich and diisopropylcarbodiimide (DIC) from Merck. Peptide synthesis grade Dimethylformamide (DMF) was purchased from Cambridge Reagents Ltd.; N-Methyl-2-pyrrolidone (NMP), triisopropylsilane (TIS), thioanisole,1,2-ethanedithiol and Dichloromethane (DCM) from Merck. Peptide grade Trifluoroacetic acid (TFA) was purchased by FluoroChem. Dithiothreitol (DTT) was purchased by AnaSpec. Reagents and solvents were used without further purification.

### Experimental Methods

#### Peptide Synthesis

Peptides were synthesized manually or on a Biotage Initiator+ Alstra via Fmoc-based synthesis with DIC/Oxyma or DIC/K-Oxyma. Typically, sidechain protected Fmoc-Amino Acids were double coupled at 4 eq. at room temperature for 45 min each. Fmoc-Thr(PO(OBzl)OH)-OH was manually double-coupled with HATU/DIEA, 2 eq. at room temperature for 40 min each. Fmoc deprotection was accomplished with 20% piperidine in DMF twice (one deprotection for 30 sec followed by an additional deprotection for 15 min). Peptides were cleaved from the resin by treatment with cleavage cocktail (95% TFA, 2.5% H_2_O, 2.5% TIS) and precipitated with diethyl ether, dissolved in 50% B, lyophilized, and subsequently purified via RP-HPLC.

#### Cloning, production and purification of recombinant proteins

Recombinant TAT-fused Apoptin has been used previously and been shown to be an effective means for protein delivery ^23^. Apoptin mutant (T106-108>A) was generated using the NEB mutagenesis kit. For semisynthesis, the C-terminus of the Apoptin encoding plasmid was removed and replaced with the Mxe-GyraseA Intein using the NEBuilder HiFi DNA Assembly Kit (New England Biolabs). Sanger sequencing of resulting plasmids was carried out by GENEWIZ.

The E.coli strain BL21-AI was used for expression of all proteins. This strain has a tightly regulated arabinose promotor in addition to the lac promotor, which therefore is good for expression of potential toxic proteins ^42^. All recombinant proteins were produced via autoinduction. A preculture was grown for 24 h at 37 °C in MDG media and used for 1:1000 inoculation into auto-induction media TYM (1% tryptone, 0.5 % yeast extract, 25 mM Na_2_HPO_4_, 25 mM KH_2_PO_4_, 50 mM NH_4_Cl, 5 mM Na_2_SO_4_, 2 mM MgSO_4_, 0.5 % glycerol, 0.05 % glucose, 0.2 % lactose, 100 μg/mL ampicillin), and grown for 16 h at 37 °C. The cells were harvested by centrifugation (4200 rpm, 30 min, 4 °C). Inclusion bodies pellets were resuspended in PBS and lysed by sonication.

10 μg/mL DNase I, 2mM PMSF and 10mM MgCl_2_ were added for 30 min at room temperature to remove DNA before the pellet was centrifuged again and resuspended in PBS with 1% Triton X-100 once more. Apoptin was solubilized from the purified inclusion bodies in 10 mL/L culture IB solubilization Buffer (25mM MES hydrate, 6M Guanidine Hydrochloride (GdmCl), 100mM NaCl and 2mM MgCl_2_, pH 6.4) by mild resuspension and incubation for 2 h at RT and centrifugation for 30 min at 30,000xg at 4 °C to remove insoluble material. Soluble material was immediately purified by nickel column and subsequent RP-HPLC. Purity of protein was assessed with analytical-HPLC and confirmed with HR-MS.

#### Reverse-Phase Chromatography

Analytical and semi-preparative reverse-phase high-performance liquid chromatography (RP-HPLC) was performed on an Agilent 1260 Infinity II instrument equipped with a dual wavelength UV-VIS detector. For analytical work, a C3 (proteins) or C18 (peptides) Zorbax column (5 μm, 4.6 × 150 mm) a were used at a constant flow of 1 mL/min. Preparative purification was performed on an Agilent 1260 Preparative HPLC system using a reverse phase Zorbax with C3 (proteins) or C18 (peptides) Zorbax column (5 μm; 21.2 × 100 mm) at a flow rate of 20 mL/min. All runs used 0.1% TFA (trifluoroacetic acid) in water (solvent A) and 0.1% TFA in acetonitrile (solvent B).

Typical gradients are described below:

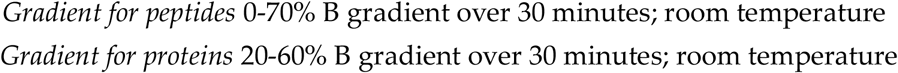

#### Mass Spectrometry

High resolution mass spectrometry was performed on a Waters Xevo G2-XS QTof. Proteins were analysed on a Waters Acquity UPLC Protein BEH C4 column (300A, 1.7 μm, 2.1Ö50 mm). Peptides were analysed on a Waters Acquity UPLC BEH C18 column (1.7 μm, 2.1×50mm) with constant flow of 0.2 mL/min using a gradient of water containing 0.1% formic acid (solvent C) and acetonitrile containing 0.1% formic acid (solvent D).

Typical gradients are described below:

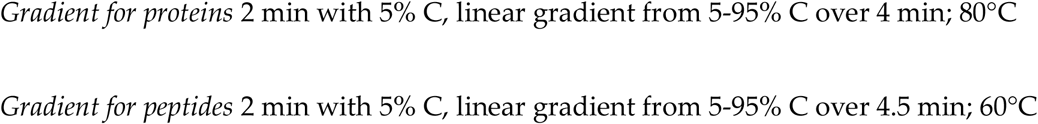

#### Intein thiolysis

For intein cleavage, Apoptin-Intein was re-dissolved in IB solubilization buffer (∼40 mg/mL) and solubilised for 2 hours followed by pH adjustment to pH3 before insoluble material was removed. pH was neutralised and protein was then diluted in thiolysis buffer (∼5 mg/mL 25 mM Mes, 100 mM NaCl, 2 mM MgCl_2_, pH 6.5) followed by the addition of 0.1 M MESNa. The reaction was incubated at 37 °C overnight with mixing and monitored by SDS-PAGE and analytical RP-HPLC. After cleavage, the Apoptin-MESNa thioester was isolated by preparative RP-HPLC and fractions containing pure product were combined and lyophilised and the identity confirmed by MS.

#### Native Chemical Ligation

For unmodified Apoptin, lyophilised Apoptin peptide (2 mg, 3.5 mmol, 1 equivalents) and Apoptin thioester (4 mg, 1.8mmol) were dissolved in 135 μL NCL buffer (100 mM phosphate buffer containing 6 M GdmCl, degassed, pH 7) and combined with 50 mM MPAA. The reaction was incubated with shaking for 1-2 h and monitored with RP-HPLC and SDS-PAGE. Finally, the reaction was incubated with 50 mM DTT for 1 h, before purification with semi-preparative RP-HPLC. Fractions containing pure, full-length Apoptin were combined and lyophilised. The procedure was repeated at a 2x higher scale for modified T108ph Apoptin. Purification with semi-preparative RP-HPLC was repeated twice.

#### Renaturation of Apoptin

Pure lyophilised proteins were redissolved in IB solubilisation buffer to a concentration of 2 mg/mL. Determination of Apoptin concentration by HPLC was based the absorbance at 214 nm with a calculated extinction coefficient based on amino acid composition ^43^. Protein was added to dialysis tubing (10kDa cut off) and dialysed twice against renaturation buffer (200x larger volume, 25 mM MES, 100 mM NaCl, 2 mM MgCl_2_, 10 mM *β*-mercaptoethanol, pH 6.4) at 4 °C (overnight and 4 hr). Protein was removed from dialysis tubing and centrifuged for 30 minutes to remove aggregates. Renatured protein was concentrated using spin concentrators pre-treated with 10% glycerol to a final concentration around 1 mg/ml. The renaturation process was monitored with SDS-PAGE, RP-HPLC and MS.

#### Pull-down assays with biotinylated peptides

Streptavidin beads (NEB) were washed and loaded with unmodified or T108ph-modified peptides. A total of 120 μg protein from cell lysates were incubated for 1 h at 30 °C. The beads were washed thrice before proteins were eluted from the beads by boiling the samples in SDS loading buffer for 10 minutes.

#### SDS-PAGE and western blotting

Samples were resolved by SDS–PAGE using 4-15% stain-free Mini-PROTEAN TGX Gels (Biorad). For western blotting, membranes were blocked with 3% milk in TBS-T for probed using antibodies for HA (Cell signalling; 2577; Rabbit 1:1000), β-Actin (Sigma-Aldrich;A5441; Mouse 1:5000), APC1 (Cell signalling; D1E9D; Rabbit 1:1000), Exportin (Cell signalling; D6V7N; Rabbit 1:1000), Importin β1 (Cell signalling; E1F1G; Rabbit 1:1000), E-cadherin (Cell signalling; 3195; Rabbit; 1:1000). All antibodies were used according to manufacturers’ instructions. The membrane was visualised by chemiluminescence on Biorad ChemiDoc MP Imager after incubation with goat anti rabbit HRP conjugate (Biorad; 170-6515) or goat anti-mouse-HRP conjugate (Biorad; 1706516).

#### 3–(4,5-dimethylthiazol-2-yl)-2,5-diphenyltetrazolium bromide (MTT) cell viability assay

Cells were seeded in triplicate in 96-well plates (5000 cells per well) and treated with varying doses of Apoptin. After 24 hours of treatment, 20 μl of MTT (5 mg/ml, Calbiochem) solution was added to each well and incubated for 2 h, after which 150 μl of solubilisation solution (50% dimethylformamide, 0.2% glacial acetic acid, 20 mM HCl, 20% SDS) was added. After further incubation overnight the optical density was measured at a wavelength of 595 nm on a LT-4000 micro-plate reader.

#### Flow cytometry

Cells were seeded in 6-well plates and treated with 5 μg/ml Apoptin proteins. Cells were harvested after 24 h, washed twice and resuspended in ice-cold annexin V binding buffer (Thermo Fisher). Cell suspension was then incubated with annexinV (FITC) (Thermo Fisher) and propidium Iodide (PI) for 30 minutes in the dark. Samples were diluted with 400 μl binding buffer prior to data acquisition. The stained cell preparations were analyzed immediately on FACS Canto II machine (BD Biosciences) and data was analysed using FlowJo software (Tree Star Inc.). Appropriate unstained and single-stained controls were prepared for determination of compensation.

For quantification of phalloidin staining, cells were seeded in 12-well plates. Cells were then washed with PBS, fixed with 4% paraformaldehyde for 15 min, washed and permeabilized with 0.2% Triton X-100 for 15 min, washed and blocked for 30 min with 3% bovine serum albumin in TBS-tween. Cells were then incubated with phalloidin (1:50) for 30 min before analysis on FACS Canto II machine (BD Biosciences) and data was analysed using FlowJo software (Tree Star Inc.).

#### Immunofluorescence

Cells were seeded in duplicate on coverslips and allowed to attach overnight. Cells were then washed with PBS, fixed with 4% paraformaldehyde for 15 min, washed and permeabilized with 0.2% Triton X-100 for 15 min, washed and blocked for 30 min with 3% bovine serum albumin in TBS-tween. Cells were then incubated overnight at 4°C with HA (Cell signalling; 2577; Rabbit; 1:500) or E-cadherin (Cell signalling; 3195; Rabbit 1:400) antibodies. Cells were washed again and incubated for 1 hour min with secondary FITC-conjugated goat anti-rabbit antibody (Sigma-Aldrich; 1:100) and TRITC-conjugated phalloidin. Cells were then mounted in Vectashield mounting medium containing 4’,6-diamidin-2-phenylindole (DAPI). Images were acquired confocal imaging system and analysed with ImageJ.

#### Invasion Assay

Transwell inserts were coated in 300 μg/mL Matrigel (BD Biosciences) and incubated at 37°C for one hour. 2 × 10^4^ cells were treated with relevant protein and then suspended in serum-free media before being added to the top of the coated transwell. Serum-containing media was added underneath the wells to act as a chemoattractant. After 24 hours, non-invading cells were removed with a cotton swab and invading cells were fixed in 100% methanol before being stained with crystal violet. Membranes were washed, dried and mounted on microscope slides. Five representative areas of each membrane were imaged at 10x magnification and invasive capacity was calculated by totalling the number of cells counted over the 5 images per membrane.

#### Anoikis resistance assays

Cells were seeded in triplicate in poly-HEMA (1.2% poly(2-hydroxyethylmethacrylate)/95% ethanol, Sigma) coated 12 well plates (2.5 × 10^3^ cells per well). The cells were then grown for 7 days at 37°C and any spherical colonies greater than 5 cells were counted as spheres.

## Acknowledgements

This work was supported by the Wellcome Trust and the Royal Society (Sir Henry Dale Fellowship 202250/Z/16/Z to M.M.M) and the NIHR Biomedical Research Centre (Studentship to JW). We thank Sofia Margiola and Karola Gerecht for advice on protein semisynthesis; the King’s College London Chemistry Department Facility for mass spectrometry services; and Beat Fierz for valuable discussions.

## Contributions

JW, MT, MMM designed the project; JW performed and analysed experiments with guidance from MT and MMM; all authors contributed to writing the manuscript.

