## Supplementary Information for "Semisynthesis reveals Apoptin as a tumour-selective protein prodrug that causes cytoskeletal collapse"

### Supplementary Figures S1-9

**Figure S1**

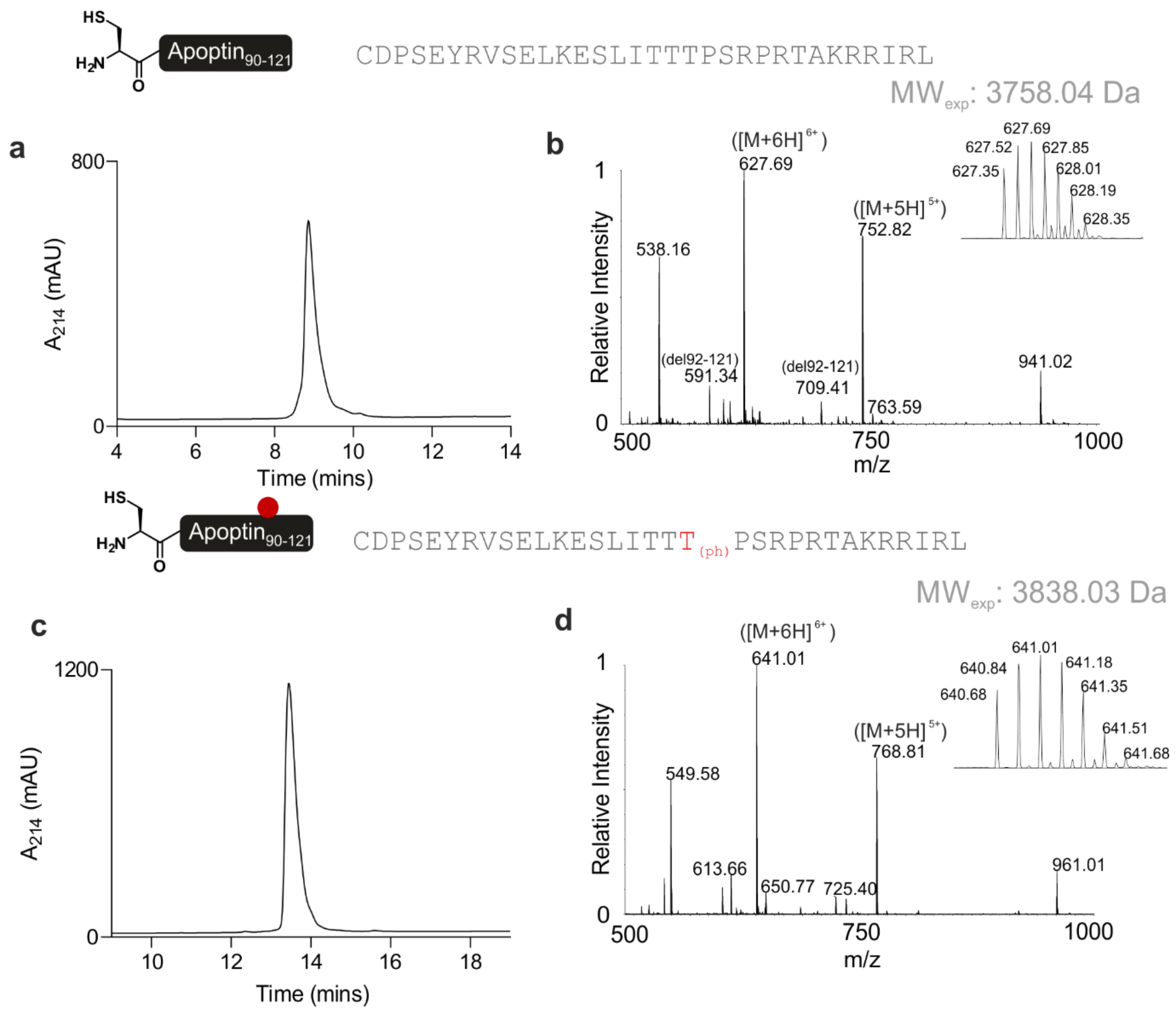

**Figure S1: Synthesis of Apoptin C-terminal peptides**

Peptides (Apoptin$_{90\text{-}121}$ and Apoptin$_{90\text{-}121}$-T108ph) were synthesis via Fmoc-SPPS and purified with RP-HPLC as described in the methods. a) RP-HPLC data and (b) MS data for the unmodified peptide (Expected mass 3758.04 Da, Observed mass 3758.06 Da). c) RP-HPLC data and (d) MS data for the T108ph peptide (Expected mass 3838.00 Da, Observed mass 3838.03 Da).

**Figure S2**

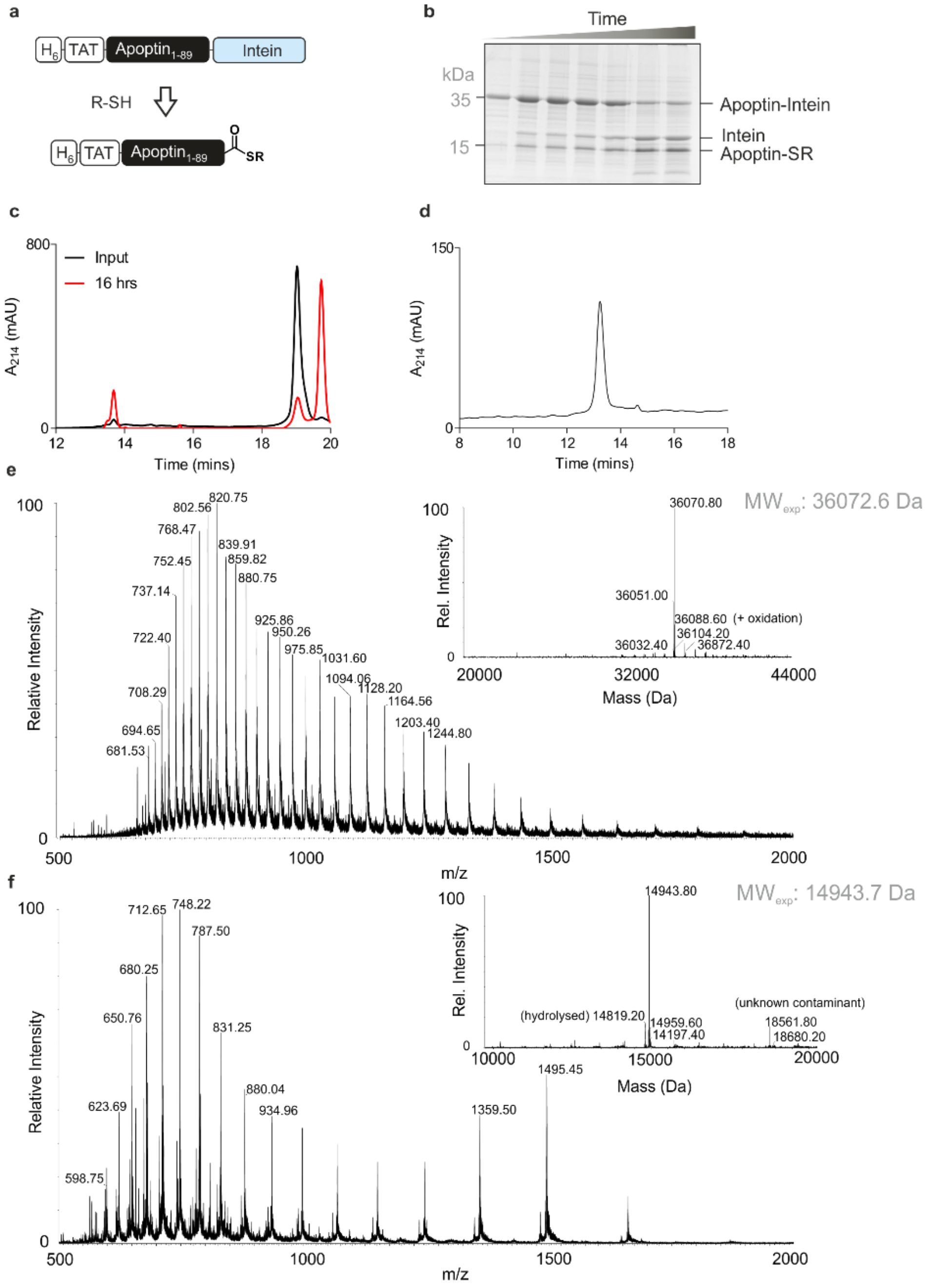

**Figure S2: Production of Apoptin-thioester**

a) Schematic for the thiolysis reaction. b) SDS-PAGE analysis of the thiolysis reaction over 16 hours. c) RP-HPLC analysis of the thiolysis reaction over 16 hours. d) RP-HPLC analysis of the isolated protein thioester. e) Mass spectrometry of the starting material corresponding to Apoptin-Intein (Expected mass 36072.6 Da, Observed mass 36070.8 Da). f) Mass spectrometry of the isolated protein thioester (Expected mass 14943.7 Da, Observed mass 14943.8 Da; hydrolysed expected mass 14819.2 Da).

**Figure S3**

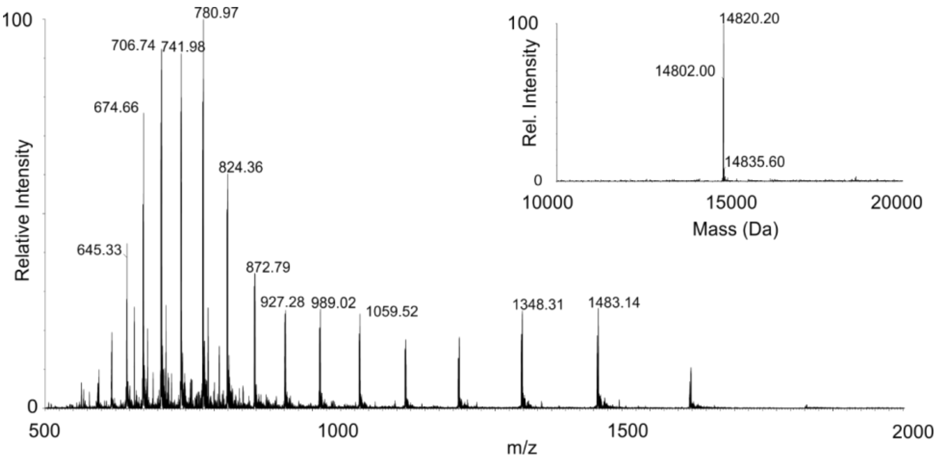

**Figure S3: Significant hydrolysis of thioester during NCL**

Mass spectrometry analysis of the thioester during the NCL reaction (Expected mass 14820.6 Da, observed mass 14820.2 Da).

**Figure S4**

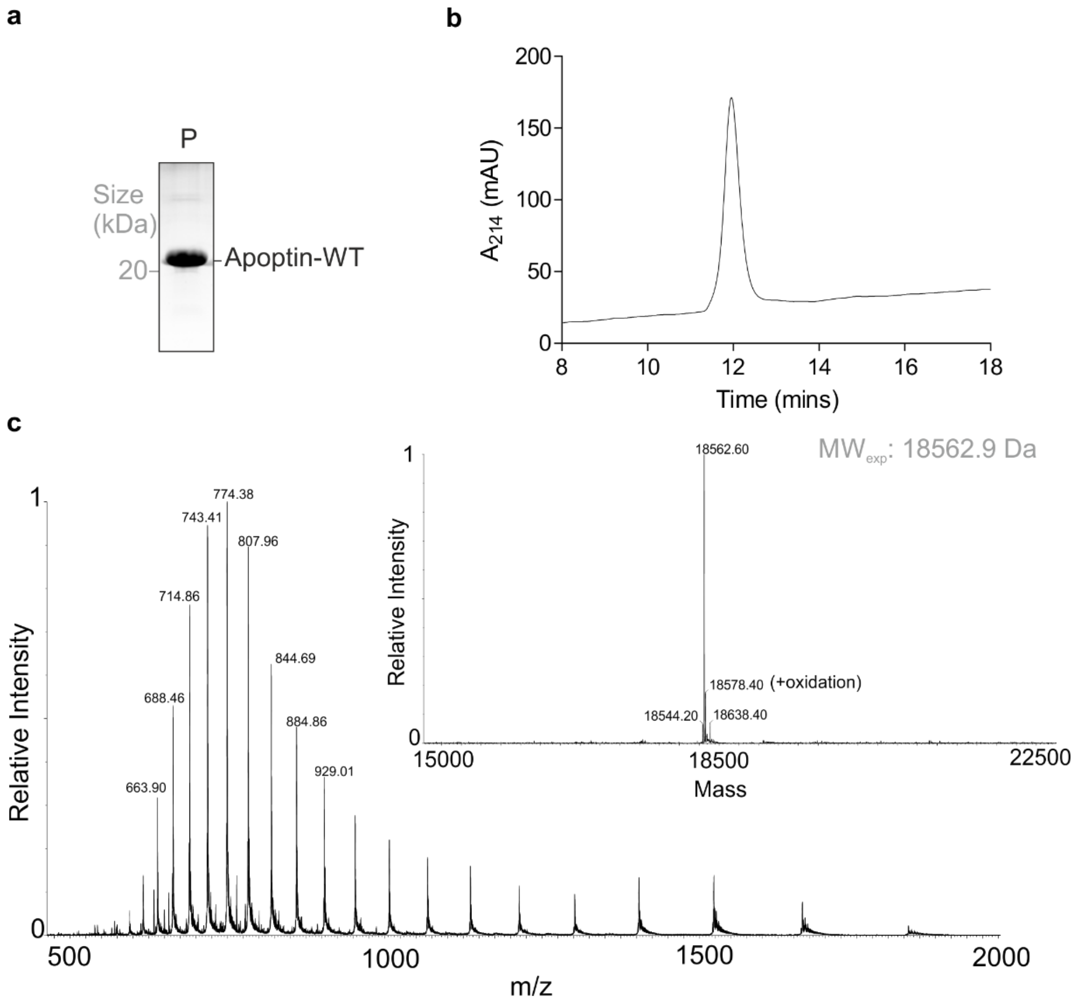

**Figure S4: Production of Apoptin-WT**

a) RP-HPLC and b) SDS-PAGE of purified and renatured Apoptin-WT. c) Mass spectrometry analysis of final product. Expected mass (-Met): 18562.9 Da, observed mass: 18562.6 Da.

### Figure S5

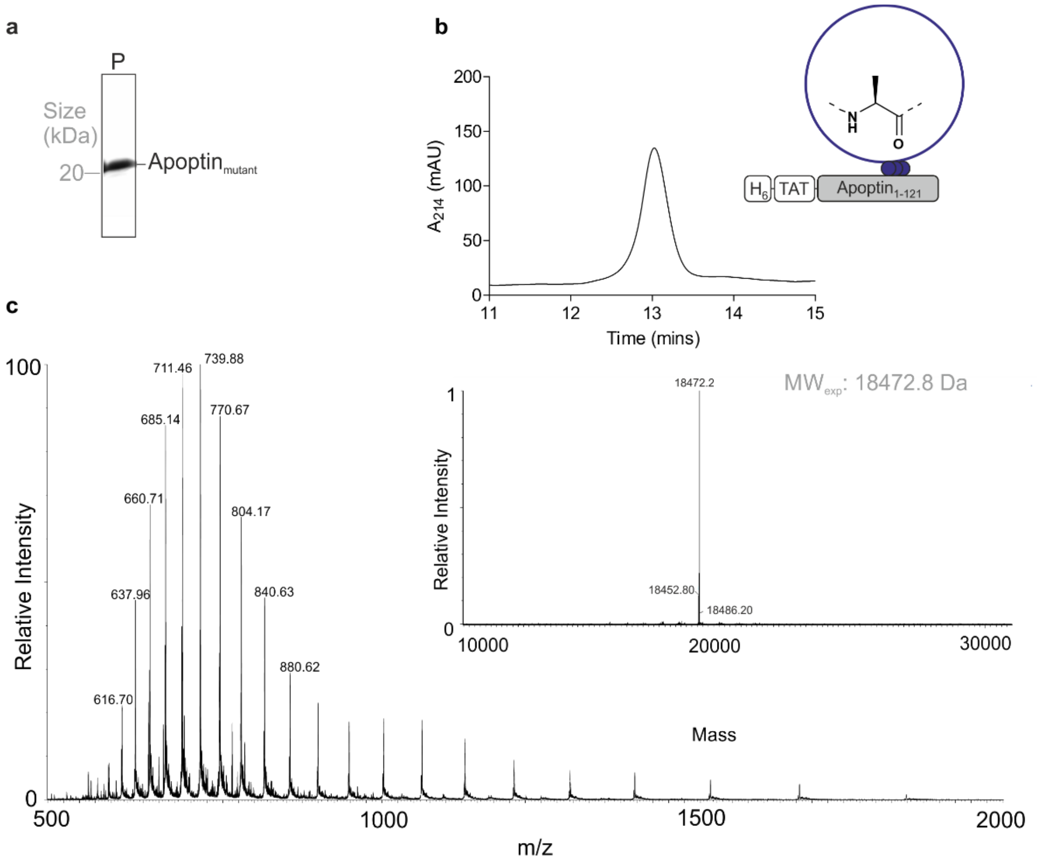

**Figure S5: Production of Apoptin-Mut**

a) RP-HPLC and b) SDS-PAGE of purified and renatured mutant Apoptin. c) Mass spectrometry analysis of final product (expected mass: 18472.8 Da, observed mass 18472.2 Da).

**Figure S6**

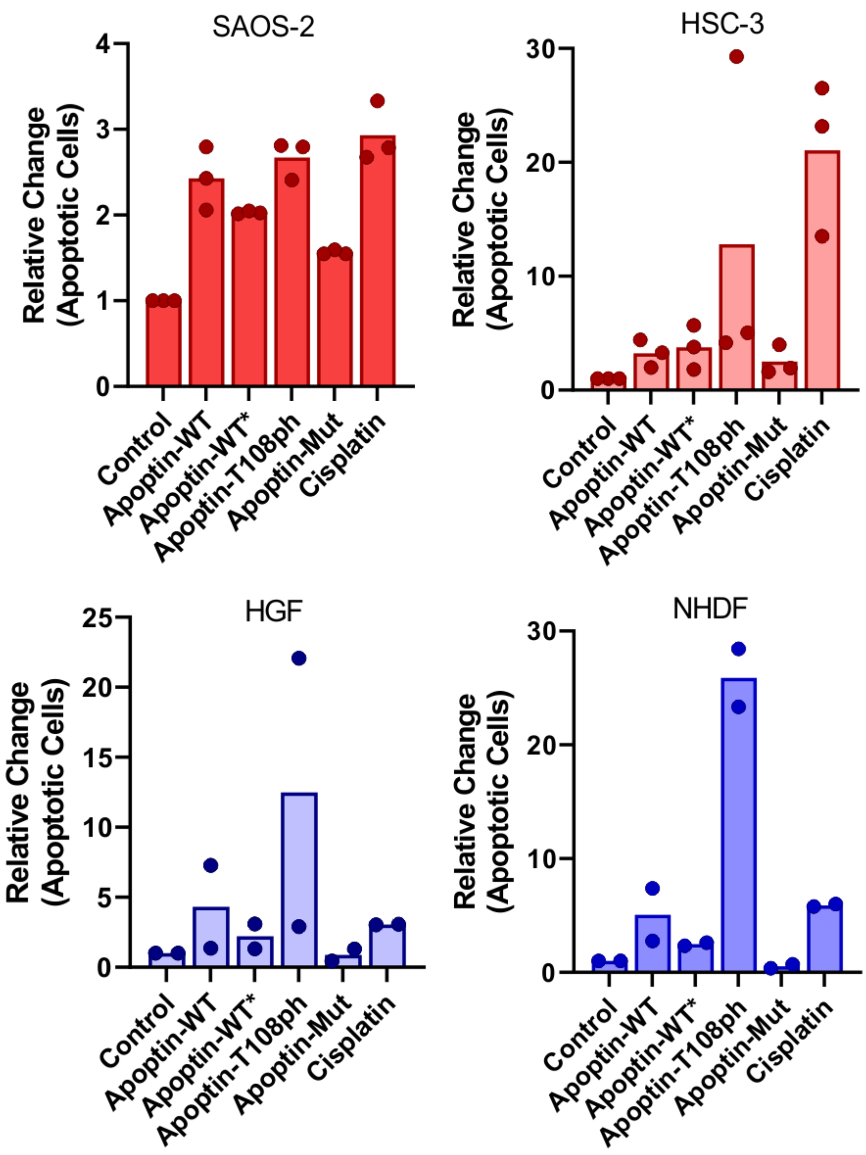

**Figure S6: Apoptotic activity of Apoptin variants analysed via FACS.**
Showing relative total Annexin V positive populations for cancer cell-lines (red, n=3) and healthy lines (blue, n=2).

**Figure S7**

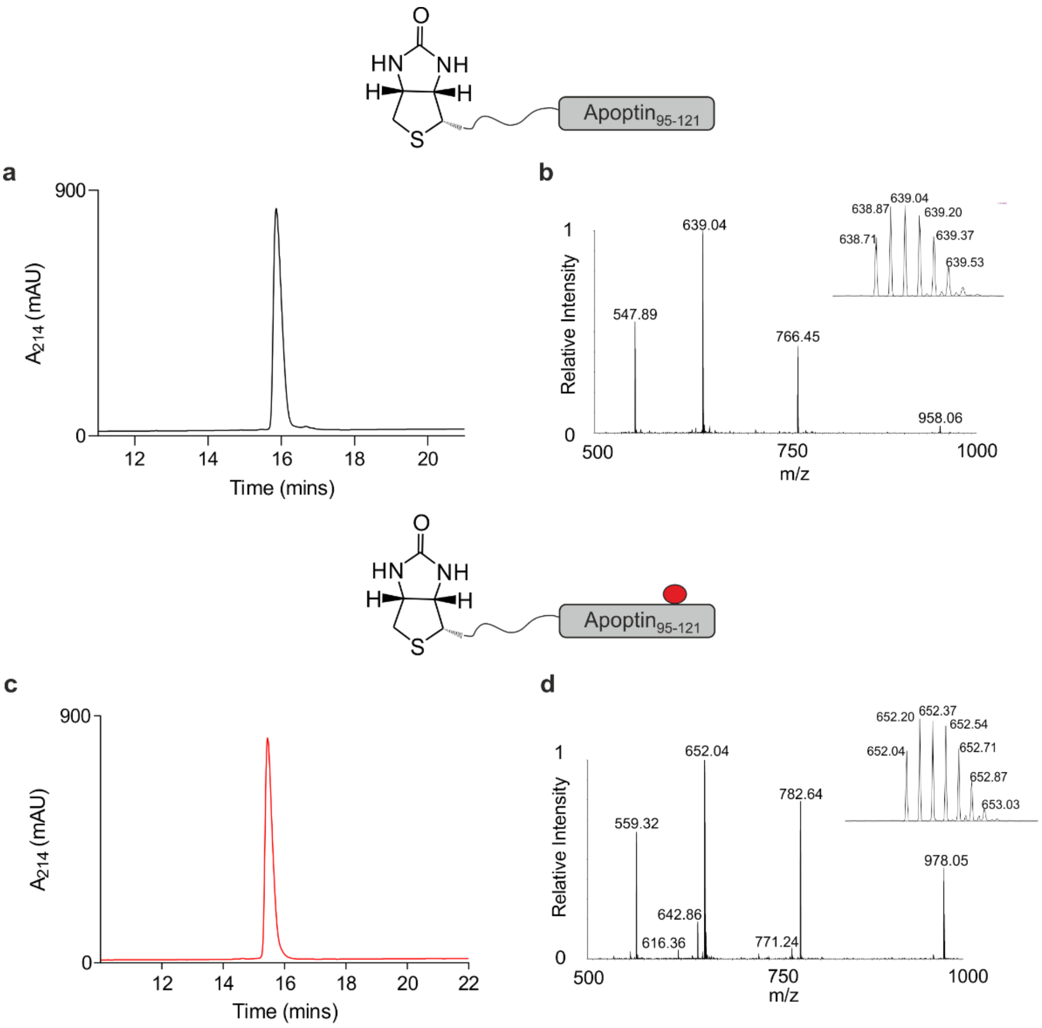

**Figure S7: Synthesis of biotinylated Apoptin peptides**

Apoptin-derived peptides were synthesis with an N-terminal biotin handle via Fmoc-SPPS and purified with RP-HPLC as described in the methods section. RP-HPLC and MS data of unmodified peptide (a, b; Expected mass: 3826.17 Da, Observed mass: 3826.26 Da) and Thr108ph peptide (c, d; Expected mass: 3906.14 Da, Observed mass: 3906.20 Da).

**Figure S8**

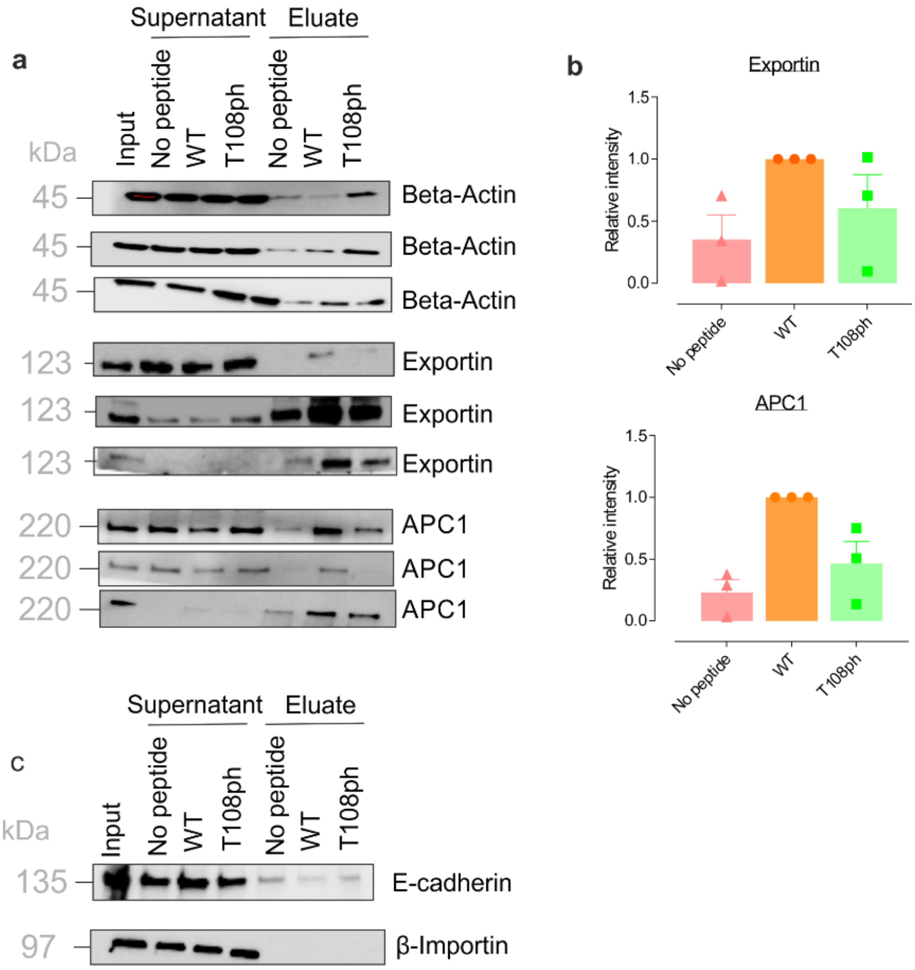

**Figure S8: Apoptin interaction to known targets**

a) Western blot analysis of beta-actin, exportin and APC1 pull-downs with wild-type and T108ph Apoptin peptides. Three independent replicates are shown. b) Quantification of Exportin (CRM-1) and APC1 pull-downs (n=3, ± SEM). c) Western blot analysis downs of E-cadherin and beta-importin pull-downs, showing that C-terminal Apoptin peptides does not interact with these targets.

**Figure S9**

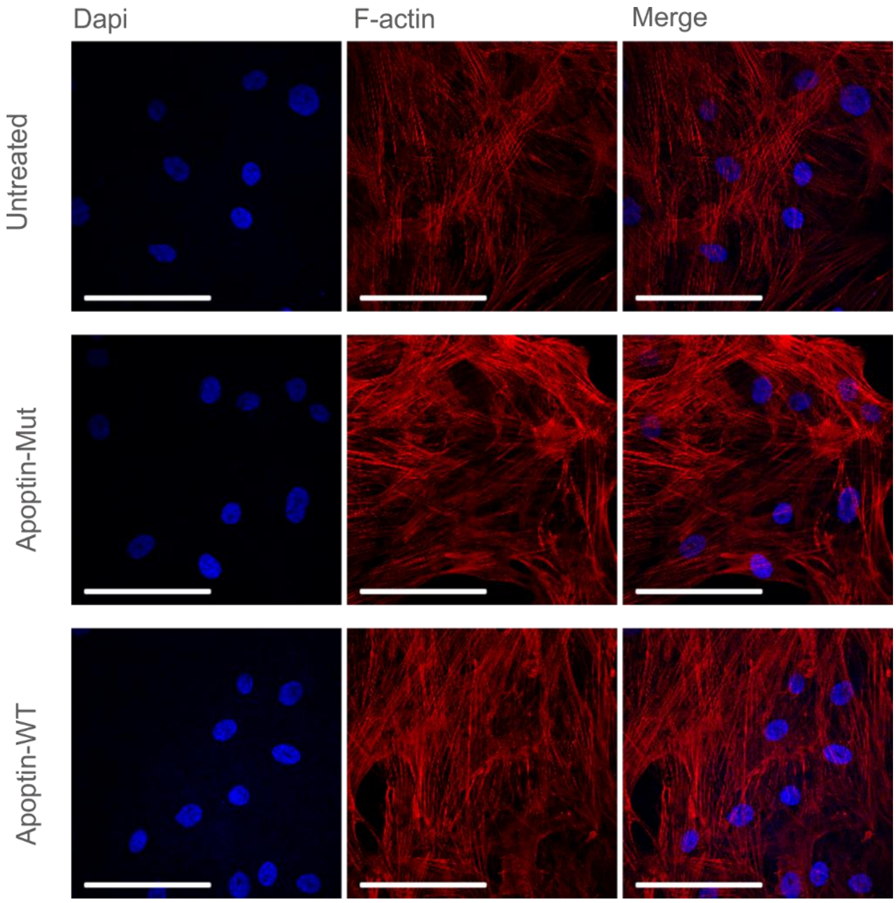

**Figure S9: Apoptin does not interfere with cytoskeleton of healthy cells**
Immuno-fluorescence images of HGF cells treated with sub-lethal doses (0.5 µg/mL) recombinant and mutant Apoptin and stained for F-Actin. Cells were imaged at 60x magnification, scale bar=40 µm.
